# Music Perception in Acquired Prosopagnosia: An Anterior Temporal Syndrome for Faces, Voices and Music

**DOI:** 10.1101/640748

**Authors:** Jason J S Barton, Jacob L Stubbs, Sebastien Paquette, Brad Duchaine, Gottfried Schlaug, Sherryse L Corrow

## Abstract

**Background:** Acquired prosopagnosia is often associated with other deficits, such as dyschromatopsia and topographagnosia, from damage to adjacent perceptual networks. A recent study showed that some subjects with developmental prosopagnosia also have congenital amusia, but problems with music perception have not been described in the acquired variant.

**Objective:** Our goal was to determine if music perception was also impaired in subjects with acquired prosopagnosia, and if so, its anatomic correlate.

**Methods:** We studied eight subjects with acquired prosopagnosia, all of whom had extensive neuropsychological and neuroimaging testing. They performed a battery of tests evaluating pitch and rhythm processing, including the Montréal Battery for the Evaluation of Amusia.

**Results:** Three of eight subjects with acquired prosopagnosia had impaired musical pitch perception while rhythm perception was spared. Two of the three also showed reduced musical memory. These three reported alterations in their emotional experience of music: one reported music anhedonia and aversion, while the remaining two had changes consistent with musicophilia. The lesions of these three subjects affected the right or bilateral temporal poles as well as the right amygdala and insula. None of the three prosopagnosic subjects with lesions limited to the inferior occipitotemporal cortex had impaired pitch perception or musical memory, or reported changes in music appreciation.

**Conclusion:** Together with the results of our previous studies of voice recognition, these findings indicate an anterior temporal agnosia syndrome that can include the amnestic variant of prosopagnosia, phonagnosia, and various alterations in music perception, including acquired amusia, reduced musical memory, and altered emotional responses to music.

---

Acquired prosopagnosia is the loss of familiarity for faces (Corrow et al., 2016b), a relatively selective problem that cannot be attributed to more general impairments in vision and memory. It is not a single disorder, but has functional and anatomic variants (Damasio et al., 1990; De Renzi et al., 1991; Barton, 2008b; Davies-Thompson et al., 2014). The lesions that are associated with acquired prosopagnosia range from occipitotemporal lesions, often involving the fusiform gyrus, to anterior temporal lesions, and are usually either bilateral or right-sided (Davies-Thompson et al., 2014), with rare subjects having only left-sided damage (Barton, 2008a).

As with all pathologic lesions, the structural damage in these subjects is often large, and often encompasses regions outside of the face network. The consequence is that prosopagnosia is frequently associated with other impairments from damage to adjacent processing circuits. Prosopagnosia from occipitotemporal lesions is often part of a ventral visual syndrome, with cerebral dyschromatopsia (Moroz et al., 2016) and topographagnosia, the inability to orient in familiar surroundings (Corrow et al., 2016a), while prosopagnosia with bilateral anterior temporal lesions can be associated with phonagnosia, the inability to recognize voices (Liu et al., 2016).

Prosopagnosia can also be developmental. Subjects with this variant do not have visible structural lesions by definition. These subjects tend to have normal colour perception (Moroz et al., 2016), though some also have problems with topographic orientation or voice recognition (Corrow et al., 2016a; Liu et al., 2015). Recently we discovered that some subjects with developmental prosopagnosia also have congenital amusia, or tone-deafness (Corrow et al., 2016c). Since amusia is neither a visual disorder nor a problem of person recognition, its co-occurrence with developmental prosopagnosia is unexpected. The basis of the association is unknown. As with the other perceptual deficits in acquired prosopagnosia, it could reflect anatomic proximity of face and music processing networks, so that they have a shared structural vulnerability to some focal disturbance. On the other hand, it may indicate that the responsible developmental failure has widespread effects on many disparate perceptual networks in vision, audition, and possibly other modalities.

If the first explanation were correct, this would predict that some subjects with acquired prosopagnosia would also have acquired amusia. The clustering of additional perceptual deficits with acquired prosopagnosia almost always reflects the close physical relationship between different perceptual networks within ventral occipitotemporal cortex. Although neurodegenerative pathology is somewhat diffuse, a supportive observation is one report of impaired familiarity for both faces and melodies in frontotemporal dementia (Hsieh et al., 2011). In this study, we examined eight subjects with acquired prosopagnosia with tests of the perception of pitch, rhythm and musical memory. Our goal was to determine if any showed music impairments, and if so, what type of impairment. We then reviewed their neuroimaging to determine whether acquired impairments of music perception were associated with a specific anatomic variant of prosopagnosia.

## METHODS

### Subjects

All prosopagnosic and control subjects could hear well enough to converse comfortably with the experimenter and none reported a hearing impairment. All subjects were fluent in English, had lived in Canada or the United States for a minimum of 10 years, most of them having spent the majority of their lives in Canada. The institutional review boards of the University of British Columbia and Vancouver Hospital approved the protocol, and all subjects gave informed consent in accordance with the principles of the Declaration of Helsinki.

Eight subjects with acquired prosopagnosia were recruited from the website www.faceblind.org or from a local neuro-ophthalmologic clinic (Table 1). These were 2 female and 6 male subjects with a mean age of 41.5 years (s.d. 15.6, range 23 to 62 years). These subjects were part of a cohort that has been studied extensively, whose details are also given in other recent reports (Hills et al., 2015; Moroz et al., 2016; Liu et al., 2016; Corrow et al., 2016a). All subjects had a neuro-ophthalmological history and examination, including Goldmann perimetry. All had corrected Snellen visual acuity of at least 20/30 in the better eye. All complained of impaired face recognition in daily life and none had complaints of mistaking one type of object for another.

**Table 1.**
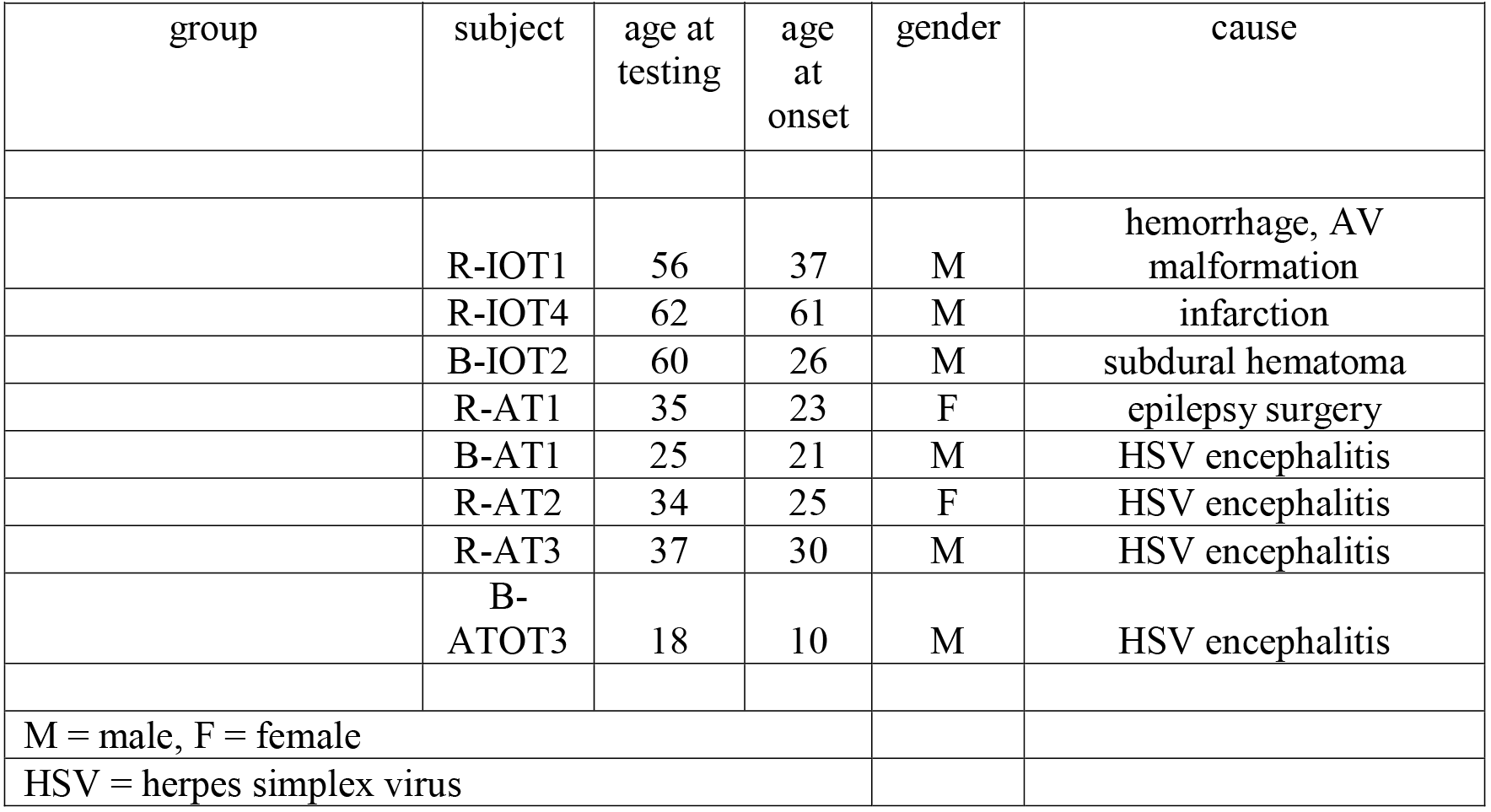
patient data.

Subjects underwent a battery of neuropsychological tests for handedness, general intelligence, executive function, memory, attention, visual perception and language skills (Table 2). The diagnosis of prosopagnosia was supported by performance on face recognition tests (Table 3). Subjects were impaired on at least one of two tests of familiarity for recently viewed faces, the Cambridge Face Memory Test (Duchaine et al., 2006) or the face component of the Warrington Recognition Memory Test (Warrington, 1984), while performing normally on the word component of the latter. Face recognition was evaluated with a Famous Faces Test (Barton et al., 2001). Although not part of the diagnostic criteria for prosopagnosia, subjects were also assessed for perceptual discrimination of faces with the Benton Face Recognition Test (Benton et al., 1972), the Cambridge Face Perception Test (Duchaine et al., 2007), and a face imagery test (Barton et al., 2003). Six of these subjects had also participated in a prior study of the discrimination of and short-term familiarity for voices (Liu et al., 2016): these results are listed in Table 3.

**Table 2.**
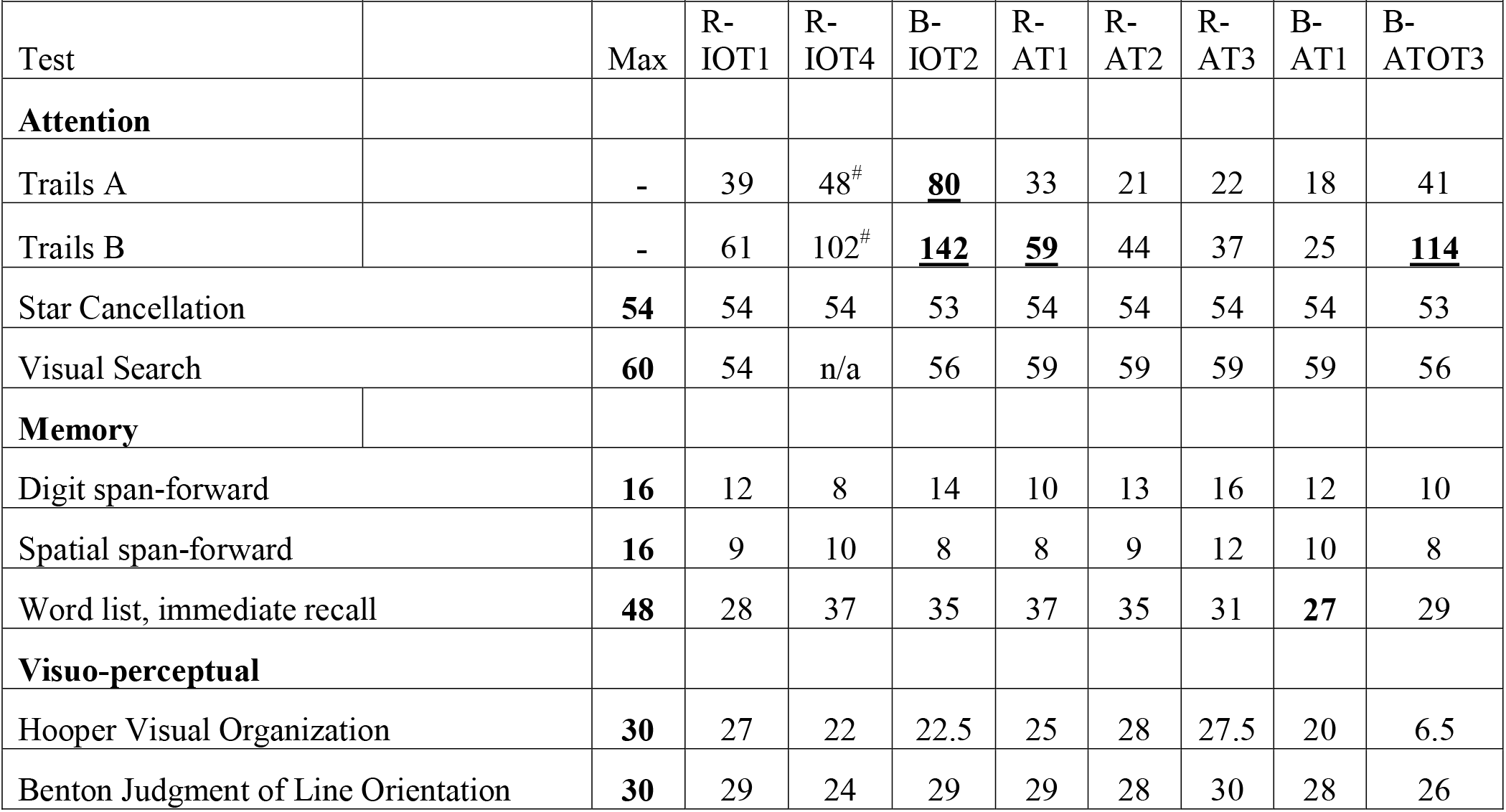
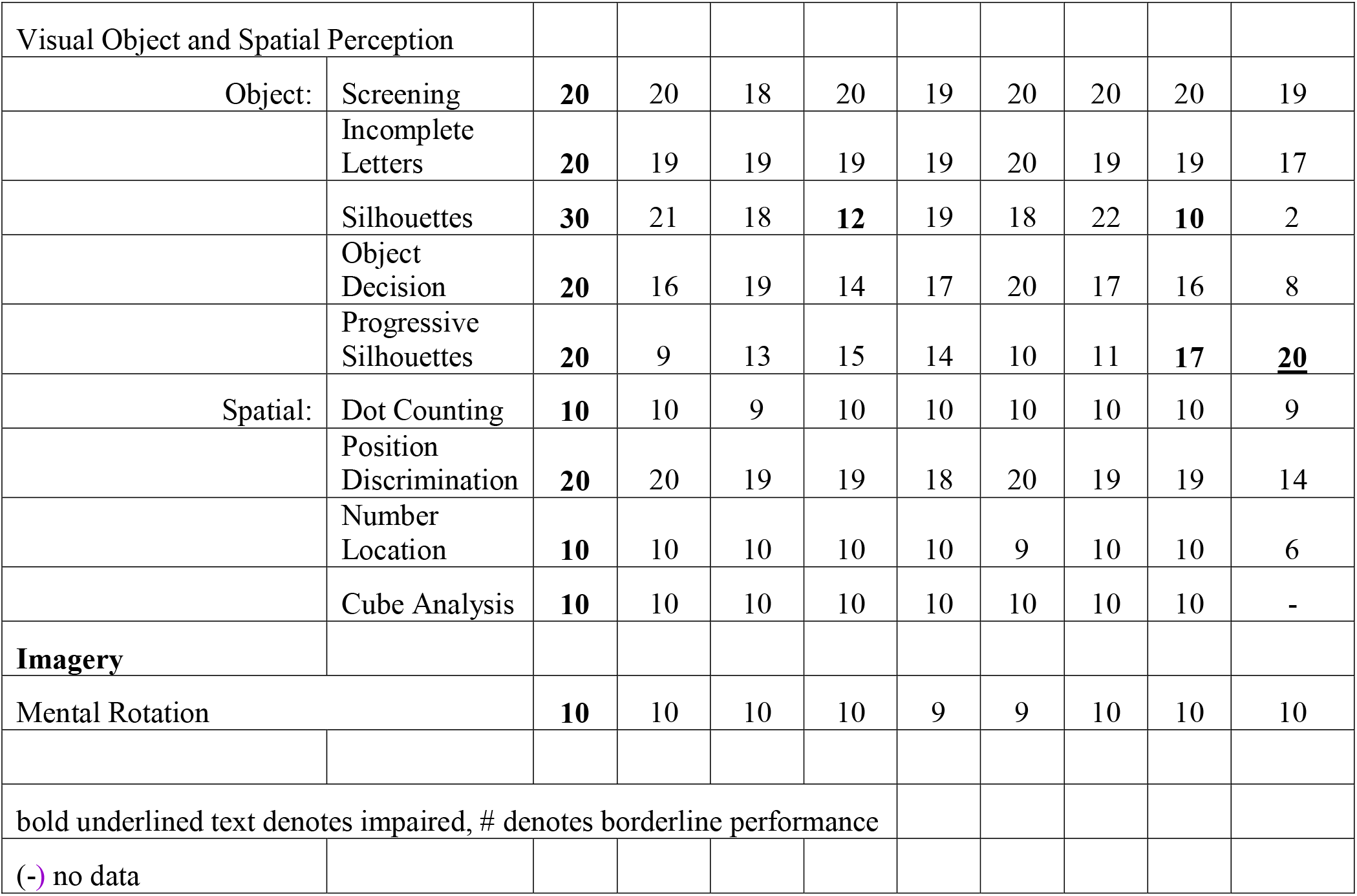
Neuropsychologic test results.

**Table 3.**
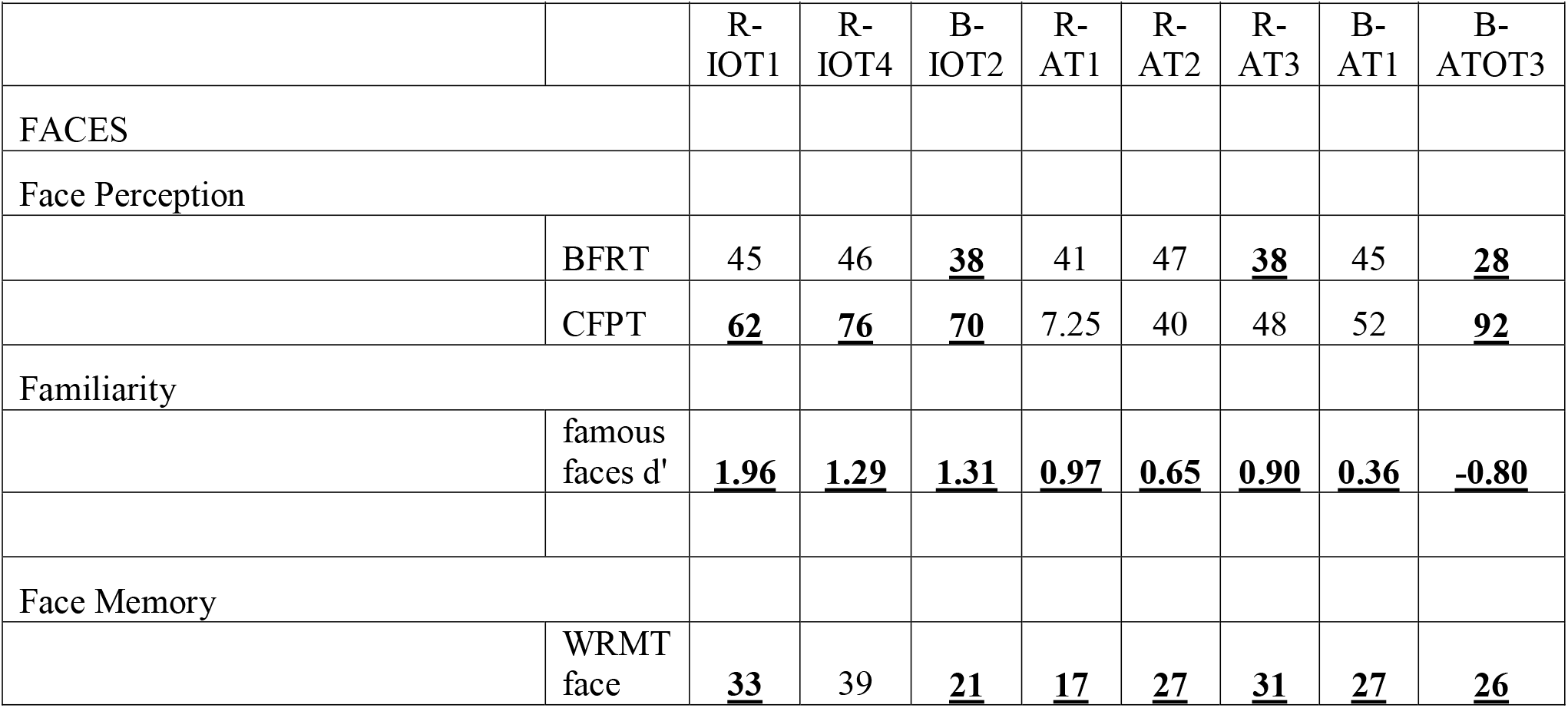
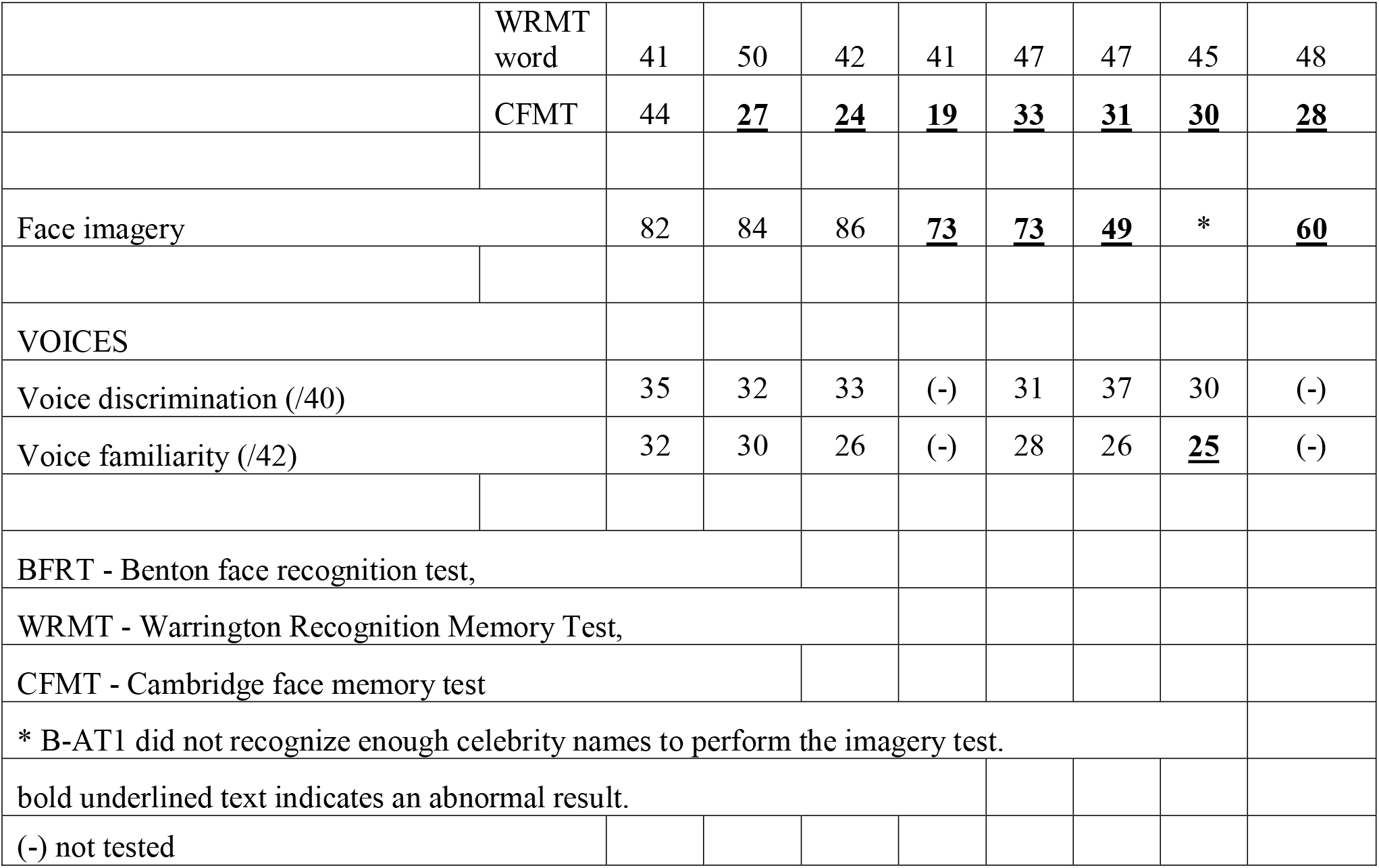
Results for Face and Voice Processing.

All prosopagnosic subjects had structural magnetic resonance imaging (Figure 1) as described in recent reports about this cohort (Liu et al., 2016; Pancaroglu et al., 2016; Hills et al., 2015). The nomenclature for our prosopagnosic subjects follows primarily the tissue loss or hypointensity on T1-weighted images. Lesions mainly anterior to the anterior tip of the middle fusiform sulcus (Weiner et al., 2014) were designated as anterior temporal (AT) and those posterior to it as inferior occipitotemporal (IOT). B-OTAT3 had a more complex lesion, with extensive right-sided lesions from the occipitotemporal to anterior temporal and parietal regions, with FLAIR sequences also showing white matter hyperintensity in the left anterior temporal and occipito-temporo-parietal regions.

**Figure 1.**
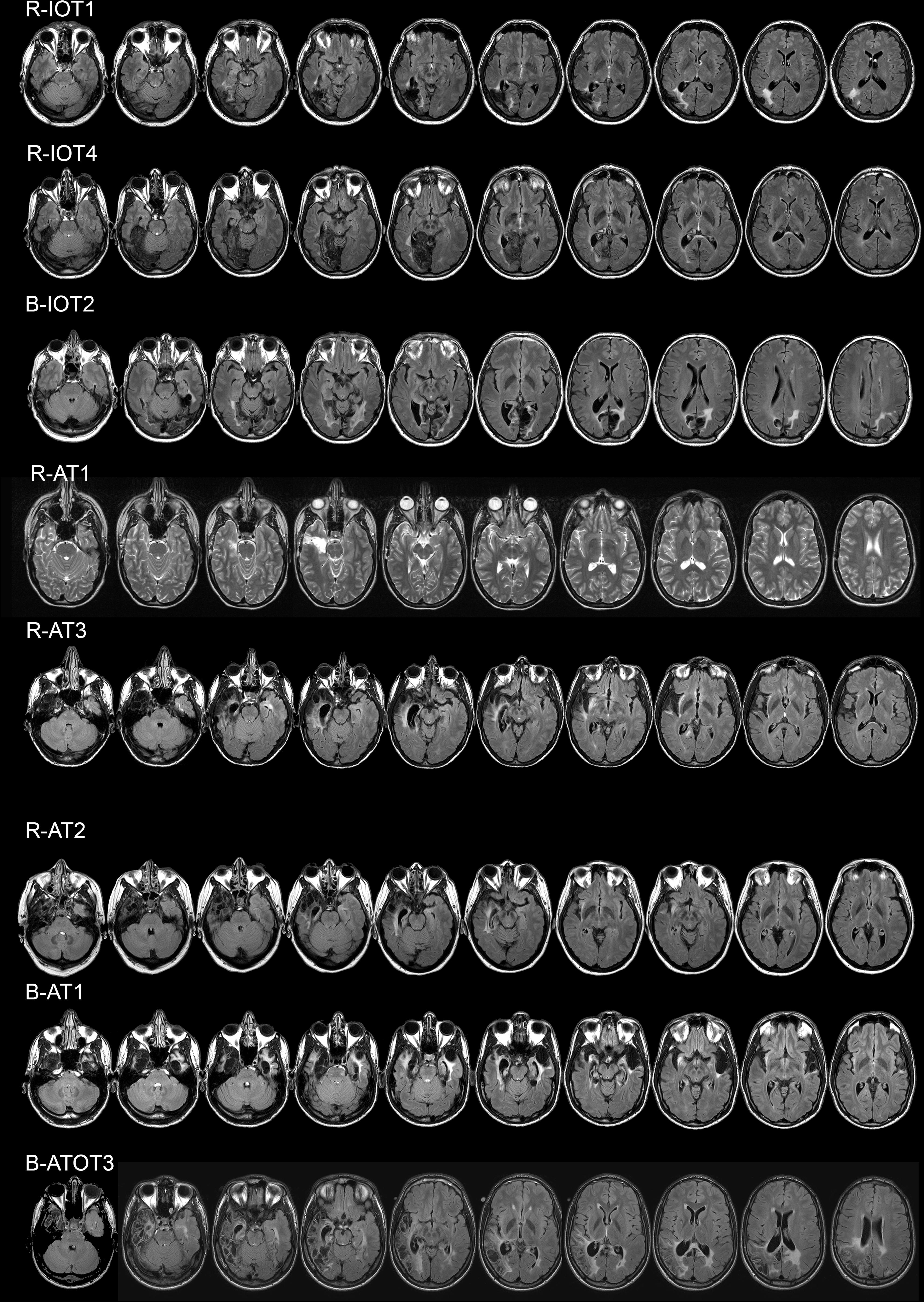
Axial MRI scans for the eight subjects with acquired prosopagnosia. FLAIR images are shown for all except for the T2-weighted image for R-AT1.

Tests of music perception were also performed by 24 control subjects (16 female, mean age 45.0 years, range 24 – 70). Subjects were excluded if they reported a history of a neurologic disorder or had best-corrected visual acuity of worse than 20/60 in their best eye. To be considered, control subjects had to affirm that they did not believe that they had any trouble recognizing faces. To guard against undiagnosed developmental prosopagnosia or amusia in our initial sample of 27 control subjects, we excluded two subjects for having a score on the Cambridge Face Memory Test of less than 44 and one for scores below published criteria on both the Montréal Battery of Evaluation of Amusia (Peretz et al., 2003) and Distorted Tunes Test (Pfeifer et al., 2015; Kalmus et al., 1980).

### Music assessments

We administered several tests of pitch and rhythm perception to confirm that any abnormality was consistent across different methods of testing. All tests were hosted online, and detailed instructions were emailed to subjects. Subjects were instructed to adjust their headphone volume to a comfortable level during practice stimuli before completing the test battery.

The *Montréal Battery for Evaluation of Amusia* (http://musicianbrain.com/mbea/) (Peretz et al., 2003) is the current “gold standard” for assessing deficits in music perception (Wilcox et al., 2015) and is the most frequently used test for diagnosing congenital amusia (Vuvan et al., 2015; Pfeifer et al., 2015). We administered all six sub-tests, each evaluating different aspects of music perception, including scale, contour, interval, rhythm, meter, and incidental musical memory recognition. Each test begins with written instructions and examples. The first three tests evaluate scale, contour, and interval components of pitch perception. In each, subjects hear two melodies sequentially and indicate if the two musical phrases are the same or different. In the fourth test the two phrases differ only in rhythm, not pitch. The fifth test, of meter, presents a single melody and subjects indicate if the melody was a waltz or a march. Finally, the sixth test examines short-term incidental memory for music: half the trials were musical phrases that had been heard in the first five tests, while half were new phrases, and subjects respond whether or not they had heard the phrases before. The scale, contour, interval, and rhythm tests contain 30 trials each, plus one catch trial in each test, which was subsequently removed. The meter and memory tests contain 30 trials. Scores for each test are calculated as the number correct out of 30. As in previous studies (Peretz et al., 2009; Gosselin et al., 2015), we calculated a subject’s composite melodic score as the average of their scores for the first three subtests (scale, contour, and interval) with pitch components. Occasionally, a participant failed to provide an answer for a trial. This occurred on a total of 7 trials for the control group, and 3 trials for the prosopagnosic group. In all, no more than one trial was missing for a test. To provide a conservative estimate of performance in these cases, missing trials were counted as correct.

The *Distorted Tunes Test* was originally described by Kalmus et al. (1980) and later updated and made available online (https://www.nidcd.nih.gov/tunestest/take-distorted-tunes-test)(Drayna et al., 2001). Twenty-six melodies of well-known North American tunes are played on a piano. Nine are correctly played but seventeen contain a wrong note, off by up to two semi-tones of the correct note, but still following the contour of the original melody. Subjects respond whether the melody was played correctly. Scores are reported as the number correct out of 26.

The *Pitch Discrimination Test* (http://musicianbrain.com/pitchtest/) (Loui et al., 2009) presents two tones sequentially, and subjects indicate if the second tone was higher or lower in pitch than the first. This test uses a staircase design, and subjects continue the task until they complete six reversals in the staircase: the average of the values at these 6 reversals is their pitch discrimination threshold, expressed in Hz. Subjects completed the task twice, and their final threshold estimate was the average of the two. This test complements the prior two tests by evaluating fine pitch discrimination outside of a musical context, with less demands on attention and memory.

The *Harvard Beat Assessment Test* (http://musicianbrain.com/hbat/) (Fujii et al., 2013) complemented the two subtests for rhythm and meter in the Montréal Battery of Evaluation of Amusia, as those subtests are useful but not infallible measures of temporal aspects of music perception, since alternative strategies can be used by participants with rhythm processing deficits (Tranchant et al., 2015). This task uses a computerized version of the Beat Finding and Interval Test described in Fujii et al. (2013). It has two components; one for beat perception and one for beat production, each using a staircase design. Both components have a repeating rhythm tapped out on a woodblock. This rhythm consists of one quarter-note, two eighth-notes, one dotted-quarter-note, and one eighth-note. In the beat perception component, subjects indicate by a key press whether the beat is accelerating or decelerating across its repetitions. In the beat production component, subjects listen to this woodblock rhythm and tap the space bar to the “beat” of the rhythm in quarter-notes. The rhythm accelerates or decelerates, and the test determines if subjects made a corresponding increase or decrease in tapping frequency. An adaptive two-alternative forced-choice discrimination paradigm was used to advance the test. A parameter was halved when the pattern of the stimulus matched with the participant’s response twice consecutively, but doubled otherwise. Every time the direction of parameter change reversed from down to up or from up to down, the parameter at which this occurred was recorded as an inflection point. One run of this task continued until six inflection points were collected. The average across these six inflection points was defined as the perception/production threshold.

Several months after completion of testing, we asked our prosopagnosic subjects two simple written questions by email, first whether they were aware of any problems with music perception or tone-deafness prior to the lesion causing prosopagnosia, and second if their music perception had changed after the onset of prosopagnosia.

### Analysis

Because our subjects span a wide age range, and there is evidence of age-related declines in music perception and performance on tests such as the Montréal Battery for Evaluation of Amusia (Moreno-Gomez et al., 2017), we determined if performance correlated with age, in particularly focusing on pitch-related tasks.

We then determined if any subject met a standard published criterion for amusia (Pfeifer et al., 2015; Chen et al., 2015; Marin et al., 2015), namely a summed score of 65 or less on the 3 melodic subtests – i.e., scale, contour and interval – of the Montréal Battery of Evaluation of Amusia, which equals the sum of the cut-off scores reported in Peretz et al. (2003).

To integrate the results of the other tests, we next derived for each subject a global score for pitch perception, by averaging the z-scores across relevant tasks from the different tests. This included the Pitch discrimination test, the Scale, Contour, and Interval subtests of the Montréal Battery for Evaluation of Amusia, and the Distorted Tunes Test, for a total of 5 tests. At an individual subject level, to determine how many subjects with prosopagnosia had impaired pitch discrimination, we calculated 95% prediction intervals from the control data (Whitmore, 1986) and classified each subject as normal or impaired by this criterion.

Similarly, we obtained a global score for rhythm perception, including the results from the Rhythm and Meter subtests of the Montréal Battery of Evaluation of Amusia, and the two tests of the Harvard Beat Assessment Test. We again calculated 95% prediction intervals for classification of the performance of individual subjects.

In the control group, global pitch and global rhythm scores were highly correlated (r = 0.68, p < 0.001). To determine if any prosopagnosic subject had a deficit greater for pitch than for rhythm perception, we regressed out the variance related to rhythm perception and used the residual variance to calculate 95% prediction intervals appropriate for single-subject comparisons for the relationship between pitch and rhythm scores.

Third, we reviewed the test assessing short-term incidental memory for music. This was the sixth test of the Montréal Battery for the Evaluation of Amusia. Because of the small number of trials, we compared the results to published normative criteria from large samples of 100 or more subjects (Peretz et al., 2003; Nan et al., 2010; Cuddy et al., 2005), which indicate that a score of 21 or less is abnormal.

## RESULTS

In this sample of healthy subjects we did not find an age-related decline of either the global pitch score (r = 0.32, p = 0.12) or the global rhythm score (r = 0.04, p = 0.86). The correlation of age with the total score for the Montréal Battery for Evaluation of Amusia also did not reach significance (r = 0.29, p = 0.16). Looking at specific pitch-related scores, we did find an age-related decline in the pitch discrimination test (r= 0.41, p < 0.038) but not for the Distorted Tunes Test (r = 0.08, p = 0.70) or the summed score for the scale, contour and interval subtests Montréal Battery for Evaluation of Amusia (r = 0.26, p = 0.22).

Four subjects of our eight prosopagnosic subjects met a standard criterion for amusia, scoring 65 or less for the sum of the performances on the scale, contour and interval subtests of the Montréal Battery for Evaluation of Amusia (Table 4). These were R-AT2 (score of 51), R-AT3 (score of 65), B-AT1 (score of 63), and B-OTAT3 (score of 65).

**Table 4.**
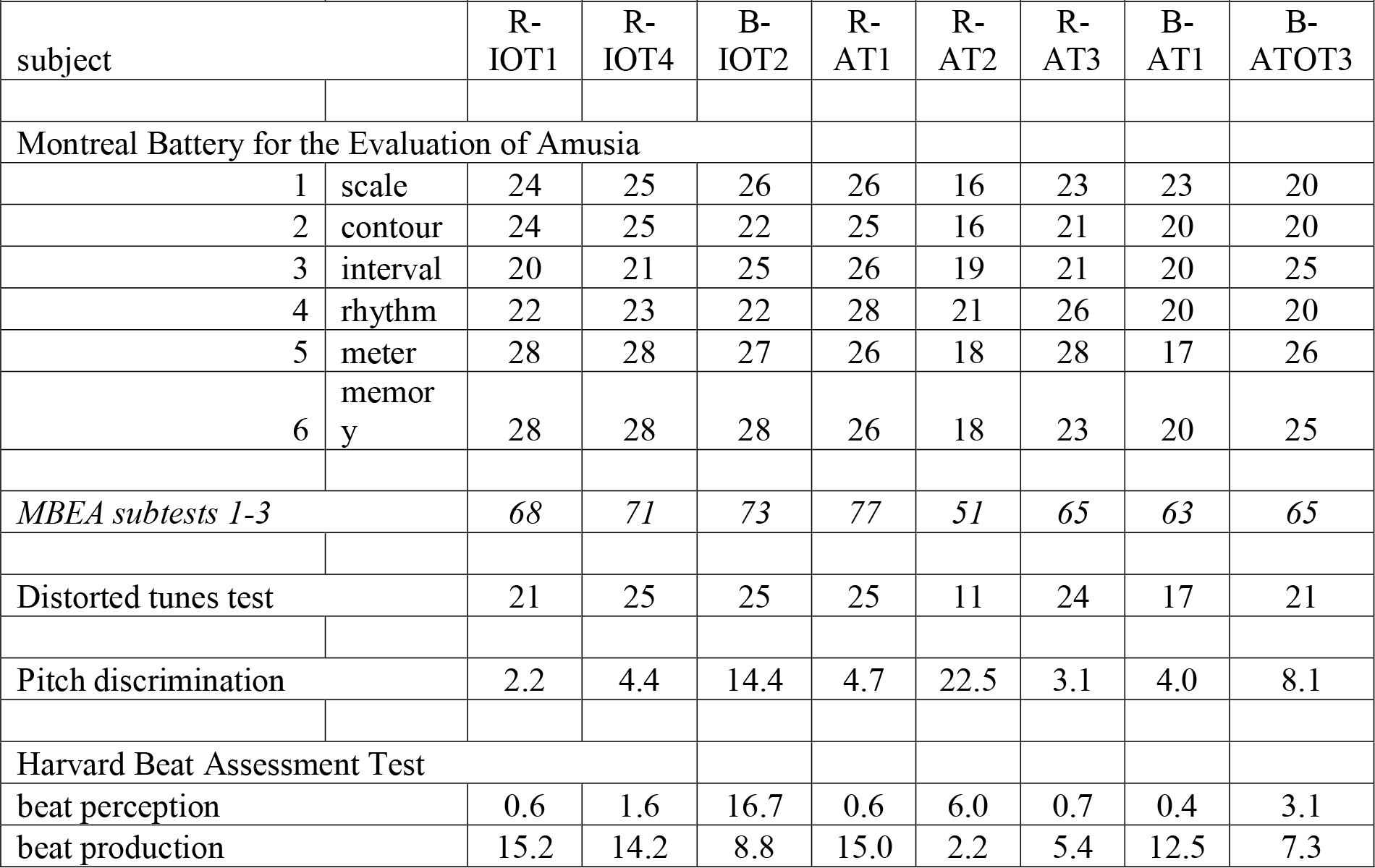
Results on music tests.

Three of these four subjects also had global pitch scores below the lower 95% prediction limit for controls (R-AT2, B-AT1, and B-OTAT3; Figure 1). For rhythm perception, only one subject, B-AT1, had a borderline global rhythm score, of −1.59. However, the analysis of the relation between pitch and rhythm scores showed that, for all three subjects with impaired pitch perception, the global pitch score was disproportionately reduced compared to their global rhythm score (Figure 2).

**Figure 2.**
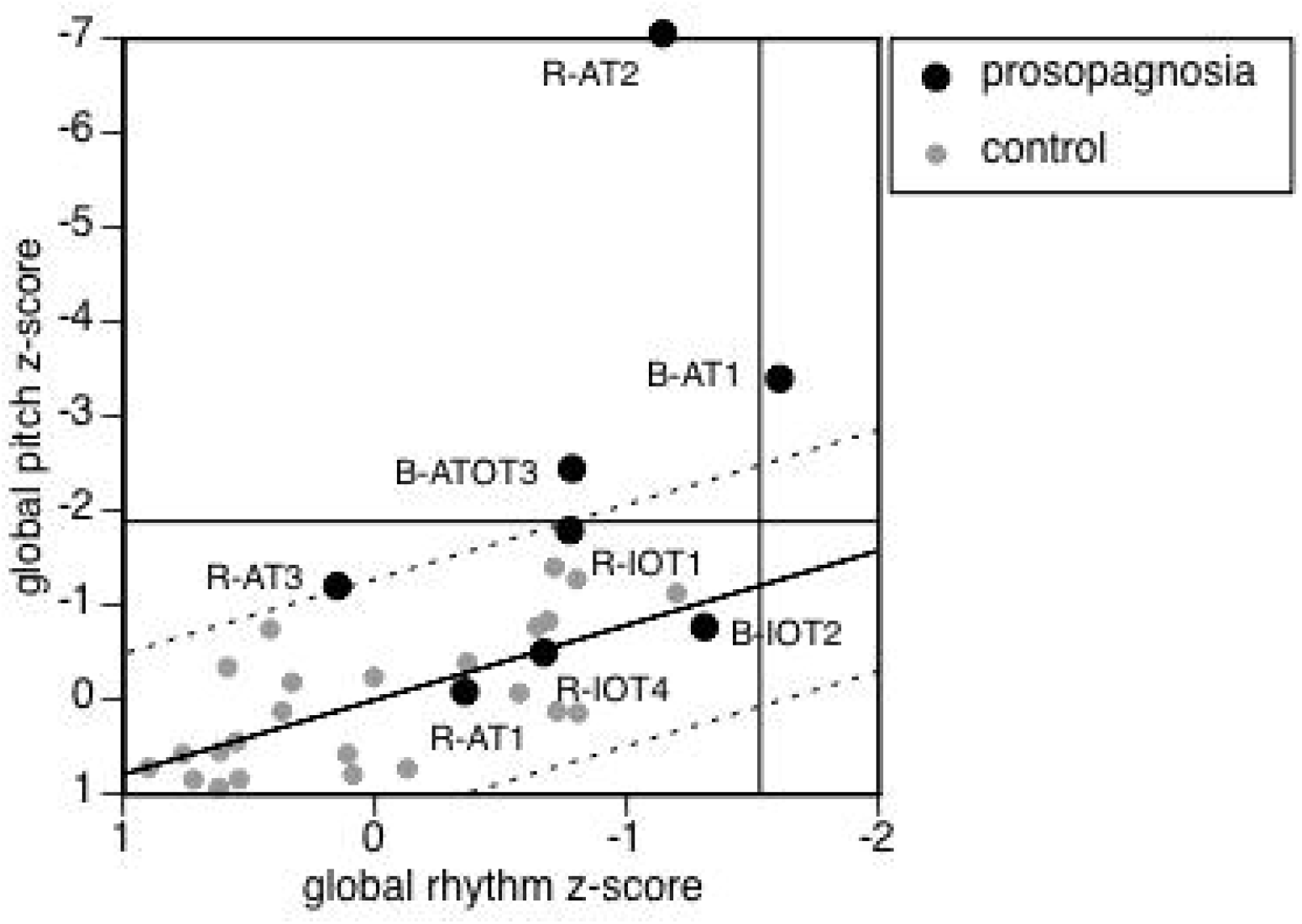
Global rhythm score plotted against global pitch score for each subject. The black solid diagonal line is the linear regression of global pitch against global rhythm scores for the control group, with the dotted black lines indicating 95% prediction limits for this regression. The horizontal and vertical solid lines indicate 95% prediction limits for global pitch and global rhythm scores respectively. R-AT2, and B-ATOT3 fall above the limit for pitch but below the limit for rhythm perception. B-AT1 also falls above the limit for pitch but is borderline for rhythm perception. However, the data for all three lie beyond the upper limit for the relationship between pitch and rhythm perception.

The global pitch score of R-AT3 was only −1.16, though he did show a borderline relative deficit when comparing his global pitch score to his global rhythm score: hence we judge that R-AT3 may have a mild deficit in pitch perception, but the evidence is not conclusive.

For short-term incidental memory for music, only B-AT1 and R-AT2 performed below the published normative criterion of 21 on the sixth subtest of the Montréal Battery for the Evaluation of Amusia (Table 4). B-OTAT3 scored 25, and of the five subjects with intact pitch perception, all scored 26 or better, with the exception of R-AT3 again, who scored 23. The Distorted Tunes Test also evaluates pitch perception in the context of familiar tunes, and thus could be considered an indirect assessment of long-term familiarity. Indeed, the two lowest scores on this test belonged to B-AT1 and R-AT2, who also had the lowest scores on the sixth subtest of the Montréal Battery for the Evaluation of Amusia (Table 4).

### Subjective impressions

None of our subjects noted a problem with music perception or tone deafness before the onset of prosopagnosia. Of the four without evidence of amusia, three reported no change after the onset of face recognition problems (R-AT1, R-IOT4, and B-IOT2). R-IOT1 reported that an unrelated life-long tinnitus had worsened and reduced his enjoyment of music. R-AT3, with the borderline deficit in pitch perception, also had not noted any change.

The three subjects with definite evidence of amusia reported alterations in musical experience after the onset of prosopagnosia. B-AT1 reported that a dislike of music was among the first changes he noted: “Beforehand, I liked music enough that I had a $3000 car stereo system with amplifiers and subwoofers. Afterwards, I went through a several year period where I preferred silence over music… For amusia, I think it’s more memory than perception”. In contrast, B-ATOT3 felt that his hearing was more acute and that he could “hear rhythms, lyrics and tones better”. He was teaching himself guitar and taking singing lessons, spending 20 hours a week on music. R-AT2 stated that her “love and appreciation for music became stronger… it seems to grab me more now and evoke lots more emotion.”

### Lesion review

R-AT2 has a lesion of the right temporal pole and amygdala, with hypertense signal in the right insula and extending posteriorly in the white matter of the temporal lobe (Figures 1, 3). B-AT1 has bilateral lesions of the temporal pole and right amygdala, and like R-AT2 had hyperintense signal in the right insula and temporal white matter. B-ATOT3 has extensive bilateral damage, which on the right includes the temporal pole, amygdala and superior temporal gyrus, with hyperintensity in the white matter underlying the right posterior insula. Of note, none of the three subjects with lesions limited to the inferior occipitotemporal cortex had amusia.

**Figure 3.**
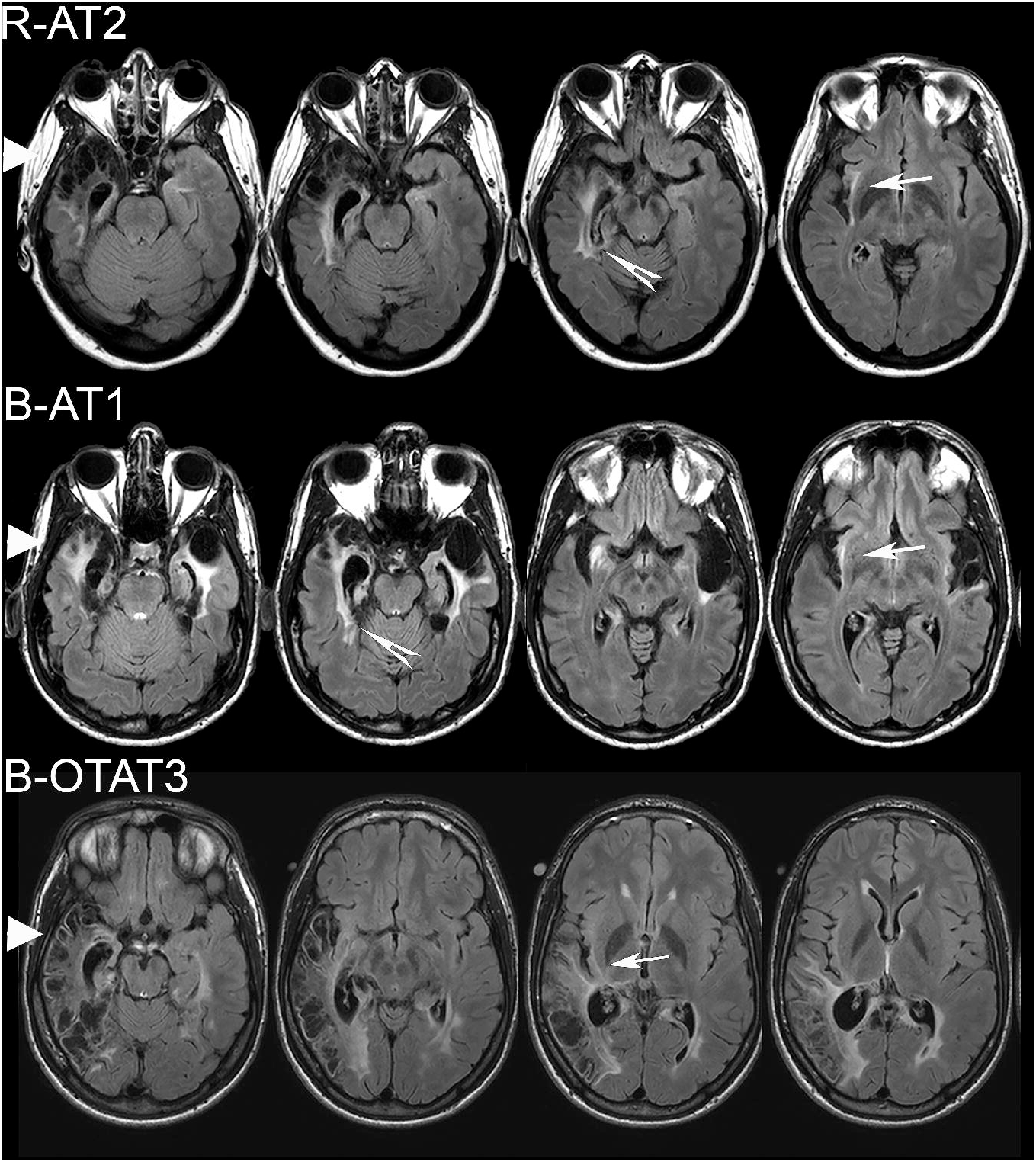
Close-up views of relevant axial FLAIR MRI sections in patients with amusia. Left-most sections show the damage to the right anterior temporal poles (triangular arrowheads) and amygdalae. White arrows with tails point to the hyperintensities in the right insulae. White arrows without tails in R-AT2 and B-AT1 point to hyperintensities in the white matter of the inferior aspect of the right superior temporal gyri. B-OTAT3 has substantial damage to the right superior temporal gyrus, as seen in right three sections.

## DISCUSSION

We found consistent evidence of impaired pitch discrimination in three of eight prosopagnosic subjects. None recalled pre-existing problems of music perception, but since none could be tested prior to the onset of prosopagnosia, and not all subjects with congenital amusia are aware of their deficit (Peretz et al., 2016), we cannot exclude the possibility that some may have had an coincidental congenital amusia. However, given an estimated prevalence of the latter in the general population of 1.5% (Peretz et al., 2016), the likelihood that a random sample of eight subjects would have three individuals with congenital amusia is less than 0.0002. Furthermore, all three with impaired pitch perception had noted a change in their musical experience. Also, though music perception declines with age (Moreno-Gomez et al., 2017), senescence was not a confound. All three of our subjects with amusia as well as the subject with a borderline deficit were under 40 years of age.

As in our subjects with developmental prosopagnosia (Corrow et al., 2016c), impaired pitch perception was a relatively specific musical deficit, as rhythm perception was generally intact. Only one subject had a borderline score for rhythm perception, and this was significantly less affected than his pitch perception. This dissociation is consistent with the results in congenital amusia (Hyde et al., 2004; Murayama et al., 2004) and most subjects with acquired amusia (Stewart et al., 2006).

We previously studied on the voice recognition of six of these subjects (Liu et al., 2016). R-AT2 performed normally on voice testing, thus providing an acquired parallel to the finding of intact voice identification in congenital amusia (Ayotte et al., 2002). B-AT1 had impaired short-term familiarity for voices, which indicates that bilateral anterior temporal lesions can result in a constellation of impairments in face, voice and music perception. His prior results also showed intact perceptual discrimination of faces and voices (Liu et al., 2016) indicating that his face and voice recognition deficits more likely reflect amnestic/associative rather than apperceptive types of dysfunction. In contrast, his amusia is characterized by an apperceptive deficit in pitch discrimination, as well as reduced memory for music and music anhedonia. Hence his problems with music and voices likely have different mechanistic origins. The co-occurrence of phonagnosia and impaired pitch perception has also been reported in two other subjects, CN and GL (Peretz et al., 1994). CN and GL had bilateral lesions involving the superior temporal gyri, temporal poles, inferior frontal gyri and insulae, similar to B-AT1.

The co-occurrence of acquired prosopagnosia and amusia suggests anatomic proximity between some components of the face and music processing networks. Neuroimaging in healthy subjects shows a right-dominant core face network in occipitotemporal regions, namely the fusiform gyrus (Kanwisher et al., 1997) and the posterior superior temporal sulcus (Haxby et al., 2000), which interacts with an extended network that includes the precuneus, inferior frontal gyrus, and anterior inferior temporal cortex (Haxby et al., 2000). Correspondingly, the neuropsychological data show that right or bilateral lesions of either occipitotemporal or anterior temporal cortex can cause prosopagnosia (Davies-Thompson et al., 2014). Occipitotemporal lesions cause an apperceptive variant, with impaired perception of facial structure, whereas anterior temporal lesions are associated with an amnestic variant, in which subjects cannot recall the appearances of faces (Damasio et al., 1990; Barton, 2008b; Davies-Thompson et al., 2014).

The music processing network also shows right hemispheric predominance for some aspects, although lateralization may be modulated by experience. Music processing depends on local connections within the temporal lobe between primary, secondary and higher order association regions, as well as projections to inferior parietal and inferior frontal regions via the arcuate fasciculus (Zatorre, 2001; Loui et al., 2009). As with faces, though, music perception is multi-faceted, and the neural substrates for pitch, timbre, melody, rhythm, memory and emotional overtones have both overlapping and distinct elements (Stewart et al., 2006). Cases of impaired pitch perception often have lesions that involve at least the anterior to middle portion of the right superior temporal gyrus and insula (Hochman et al., 2014; Terao et al., 2006; Ayotte et al., 2000; Peretz et al., 1994; Sihvonen et al., 2016), sometimes bilateral or extending to the temporal poles (Peretz et al., 1994). Our three subjects with impaired pitch perception had large right anterior temporal lesions with subtle anomalies in the right insula.

While our testing of musical memory was limited, two of these three subjects were also impaired on the incidental memory test of the Montréal Battery for the Evaluation of Amusia, and had the lowest scores on the Distorted Tunes Test, which uses familiar songs as the context for testing pitch perception. Impaired memory for pitch is seen in congenital amusia (Gosselin et al., 2009). Likewise, many acquired cases with impaired musical memory also have impaired pitch perception (Stewart et al., 2006), and hence can be considered as having an apperceptive music agnosia. This is again associated with lesions of the anterior superior temporal gyrus and insula, as in our subjects. Studies of subjects with anterior temporal lobectomy have shown impaired melodic memory after right-sided lesions and impaired memory for lyrics with left-sided lesions (Samson et al., 1991, 1992).

The narratives of our patients also suggest variable effects on the emotional response to music. This not necessarily a given, as music perception and its emotional response are dissociable: some subjects with acquired (Lechevalier et al., 1984; Peretz et al., 1998) or congenital amusia (Gosselin et al., 2015) still report enjoyment of music, while musical anhedonia can occur with intact music perception (Griffiths et al., 2004; Mas-Herrero et al., 2014). Among our cases with impaired pitch perception, B-AT1’s report is consistent with musical anhedonia and music aversion, as previously reported in some cases of impaired pitch or timbre perception from right temporal and insular lesions (Griffiths et al., 1997; Mazzucchi et al., 1982; Terao et al., 2006; Hirel et al., 2014). In a series of 73 patients with frontotemporal dementia (Fletcher et al., 2015), 14 showed music aversion, which was associated with cortical thinning in right anterior temporal cortex, entorhinal cortex, hippocampus, amygdala and bilateral insulae.

In contrast, two of our three amusic subjects reported enhanced enjoyment of and/or participation in music, consistent with musicophilia (Fletcher et al., 2015). There are case reports of musicophilia in patients with frontotemporal dementia (Geroldi et al., 2000; Boeve et al., 2001; Hailstone et al., 2009). In one this evolved in concert with right amygdala and temporal atrophy (Boeve et al., 2001). Larger series suggest that musicophilia may be present in a quarter to a third of subjects with frontotemporal dementia (Fletcher et al., 2013; Fletcher et al., 2015). Although some authors report the impression that musicophilia was often accompanied by loss of music discrimination (Fletcher et al., 2013), music perception has rarely been evaluated in musicophilia, with the exception of a demonstration of preserved memory for famous tunes in one case (Hailstone et al., 2009). Our report provides objective evidence that musicophilia can occur with impaired pitch discrimination.

The co-occurrence of acquired amusia and prosopagnosia is a novel finding. However, studies of acquired amusia generally did not include face recognition tests in their neuropsychological batteries (Griffiths et al., 1997; Peretz et al., 1994; Baird et al., 2014; Sarkamo et al., 2009a, b). One amusic study even reported on the word but not the face component of the Warrington Recognition Memory Test (Baird et al., 2014). The omission of face tests reflects the fact that the focus of their testing was on general cognitive processes of memory, attention and linguistic processing, and any testing of vision was limited to basic processes probed by the Visual Object and Space Perception battery (Baird et al., 2014; Griffiths et al., 1997; Warrington et al., 1991).

While we are not aware of prior reports of patients with both prosopagnosia and amusia specifically, some studies of frontotemporal dementia have reported co-existent problems with face and music perception. Although not directly relevant to our report, two studies established the multimodal nature of impaired emotion recognition in frontotemporal dementia by showing parallel deficits in emotional processing of music, faces and voices (Omar et al., 2011; Hsieh et al., 2012). However, recognition of facial emotions is dissociable from recognition of facial identity (Fox et al., 2011) and its impairment is not part of the definition of prosopagnosia. Also, the subjects of these two studies were not shown to have amusia: in fact, the subjects in one report performed well on the scale subtest of the Montréal Battery for the Evaluation of Amusia (Hsieh et al., 2012).

More relevant to our work are two other studies of frontotemporal dementia. One reported a patient with musicophilia and impaired face and object recognition, who performed poorly on a test of famous face recognition (Hailstone et al., 2009). The second studied 13 patients and found impaired familiarity for both famous faces and famous melodies (Hsieh et al., 2011). However, in both reports these impairments were part of more widespread semantic deficits, and neither evaluated music discrimination. Nevertheless, while these points reduce the parallel with our work, the voxel-based morphometric analysis of the latter study (Hsieh et al., 2011) did find overlap in the right anterior temporal lobe between regions implicated in impaired familiarity for famous tunes and that for famous faces, a finding that supports our conclusion that focal lesions in this region can impair both face and music processing.

A limitation of this study is the small number of patients studied; however, acquired prosopagnosia is rare and our cohort is the largest assembled in recent decades. Despite this limitation, our prior work with this group has shown that apperceptive prosopagnosia is associated with dyschromatopsia (Moroz et al., 2016) and impaired cognitive map formation (Corrow et al., 2016a) as part of a ventral visual syndrome following right or bilateral lesions of inferior occipitotemporal cortex. This accords with prior reports (Bouvier et al., 2006) but clarifies that these associations are specific to the occipitotemporal variant. The co-occurrence of these deficits is not invariable: not all patients have all three components, which is to be expected as face, colour and topographic processing involve neighbouring rather than identical perceptual networks, and the impact of a lesion in any given patient will differ across these networks. Along with our prior study of voice recognition (Liu et al., 2016), the current report points to the existence of a second syndrome, an anterior temporal agnosia syndrome, which follows right or bilateral damage and consists of the amnestic variant of prosopagnosia, phonagnosia, and various alterations of music perception, including impaired pitch discrimination, reduced musical memory, and altered emotional responses to music, such as music aversion and musicophilia. Again, not all face, voice and music deficits are present in all patients, which may reflect the degree of pre-morbid lateralization of separate processing networks as well as the anatomic extent of lesions in one or both hemispheres. Given the rich and complex nature of sensory processing, further studies will likely reveal other perceptual or recognition deficits in the same or other sensory modalities following anterior temporal damage.

## ACKNOWLEDGMENTS

This work was supported by CIHR operating grant (MOP-102567) to JB. JB was supported by a Canada Research Chair (950-228984) and the Marianne Koerner Chair in Brain Diseases. BD was supported by grants from the Economic and Social Research Council (UK) (RES-062-23-2426) and the Hitchcock Foundation. G. S. acknowledges support from NIH (DC009823, DC008796). S.P. is supported by a Postdoctoral Fellowship from the Canadian Institutes of Health Research. SC was supported by *National Eye Institute* of the National Institutes of Health under award number F32 EY023479-02. The content is solely the responsibility of the authors and does not necessarily represent the official views of the National Institutes of Health.

